# Fatty acid metabolism and dietary lipid composition shape B-cell responses

**DOI:** 10.64898/2026.10.02.756283

**Authors:** Bongwe Ngwenyama, Léa Bourguignon, Elpida Bokou Gianneli, Yannice Benazzouk, Mylène Laflamme, Isabella Giacomini, Thi Kem Nguyen, Charles Viau, Audrey L. Collin, Fabio Zani, Julianna Blagih

**Affiliations:** Centre de Recherche de l’Hôpital Maisonneuve-Rosemont, Montreal, Canada; Department of Obstetrics-Gynecology, Université de Montréal, Montreal, Canada; Department of Biomedical Sciences, Université de Montréal, Montreal, Canada; Department of Pharmacology and Therapeutics, McGill University, Montreal, Canada

**Author notes:** equal contribution (Bongwe Ngwenyama, Léa Bourguignon).

## Abstract

B-cell activation requires extensive metabolic reprogramming, yet how B cells acquire and utilize extracellular fatty acids and adapt to changes in lipid availability remains poorly understood. Here, we show that B-cell activation and the extracellular lipid environment dynamically regulate B cells. B-cell receptor (BCR) and Toll-like receptor 4 (TLR4) stimulation induced distinct mitochondrial and lipid-handling programs, differentially regulating fatty-acid uptake, cholesterol acquisition, lipid storage and mitochondrial respiration. Disrupting long-chain fatty-acid uptake, acyl-CoA activation or mitochondrial utilisation impaired B-cell proliferation, demonstrating that multiple steps of fatty-acid metabolism are required for B-cell expansion. However, increasing extracellular fatty-acid availability alone did not enhance proliferation, indicating that fatty-acid acquisition and metabolic utilisation are functionally distinct. B cells also acquired fatty acids directly from adipocytes, identifying the local microenvironment as a potential source of metabolic substrates. In vivo, dietary fatty-acid chain length differentially shaped the germinal-centre response. A long-chain fatty-acid-rich diet increased germinal-centre B cells and T follicular helper cells and induced mitochondrial and transcriptional changes that partially converged with those observed following obesogenic feeding. Circulating lipidomics further revealed shared remodelling of lipid species between long-chain fatty-acid-rich and obesogenic diets, including altered partitioning of stearate-containing lipids. Exogenous fatty acids also modified membrane order in activated B cells, suggesting that lipid availability can influence both metabolic and biophysical properties. Together, these findings establish fatty-acid uptake, activation and utilization as integral components of B-cell metabolic adaptation and demonstrate that dietary fatty-acid composition can remodel the lipid environment and germinal-centre response.

## Introduction

B lymphocytes undergo profound metabolic adaptations during activation, proliferation, and differentiation to support expansion, antibody production, and humoral immunity^1^. This metabolic plasticity is particularly important during the germinal centre (GC) response, where activated B cells undergo rapid proliferation, somatic hypermutation and affinity-based selection before differentiating into memory B cells or antibody-secreting plasma cells^2^. This metabolic reprogramming requires tight coordination between glucose and amino acids, enabling B cells to respond rapidly to antigenic stimulation and mitochondrial metabolism^3^. Key pathways involve glycolysis as well as the oxidation of the saturated fatty acid palmitate for mitochondrial ATP production^4–6^. However, despite growing appreciation of the importance of lipid metabolism in immunity, how individual key nodes of fatty acid metabolism and how dietary fatty acid species regulate primary B-cell responses remains poorly understood.

Lipids provide substrates for membrane biogenesis and mitochondrial metabolism while also acting as signalling molecules that can influence cellular differentiation^7^. Saturated fatty acids – single-bond structures – are often considered a homogeneous class of lipids, despite marked differences in their biochemical properties. Medium-chain fatty acids (MCFAs, C6-C12) and long-chain fatty acids (LCFAs, C14-20) differ substantially in cellular uptake, intracellular trafficking, mitochondrial transport and metabolic utilization^8^. These differences suggest that fatty acid chain length may have important consequences for immune cell metabolism and function. However, whether distinct saturated fatty acids differentially regulate B-cell activation, differentiation and metabolism has received little attention.

Importantly, diet profoundly influences the circulating lipid environment; dietary fat composition can remodel the circulating lipidome and alter the lipid substrates available to immune cells^9^. However, whether differences in dietary fatty acid composition influence GC B-cell responses, and how B cells metabolically adapt to fatty acids of different chain lengths, remain poorly understood. Dietary saturated fats, when consumed chronically and in excess, drive metabolic dysfunction, in part driven by B cells. Furthermore, diet-induced obesity reduces influenza antibody titers, which increased consumption of polyunsaturated fatty acids (PUFA) can rescue.^10^ However, how different dietary chain lengths of saturated fats within the advisable consumption range, or how excess saturated fat before obesity, affect B cell response remains unexplored.

In addition to diet-derived circulating lipids, adipocytes could be a major source of fatty acids in lymphoid tissues and the tumour microenvironment, given their close anatomical proximity. Through lipid release and transfer, adipocytes can actively reshape cellular metabolism in immune cells and impact their function, including in memory CD8 T cells^11^. Although B cells are present in adipose tissue and lymphoid organs are in direct contact with it, how adipocyte-derived factors regulate primary B-cell proliferation remains unknown^12^.

Here, we investigated how saturated fatty acid chain length shapes B-cell responses across systemic, cellular and metabolic scales. Using diets enriched in MCFAs or LCFAs, we examined remodelling of the circulating lipidome and the GC response *in vivo*, and subsequently defined how primary B cells acquire and metabolically adapt to fatty acids of different chain lengths. Finally, using adipocyte–B-cell co-culture systems, we investigated adipocytes as a local source of exogenous fatty acids. Together, these approaches establish a framework linking dietary fatty acid composition and the circulating lipid environment to B-cell lipid handling and humoral immune responses.

## Results

### B cells alter their mitochondrial metabolism in response to different stimuli

Immune cells rewire their metabolism depending on the external stimulus^13,14^. We measured changes in mitochondrial parameters – load and membrane potential – in response to BCR stimulation (a-IgM) and TLR activation (LPS). We found that BCR stimulation after 24 hours of activation increased both mitochondrial content and membrane potential compared to LPS stimulation and unstimulated controls (**Figure 1A**). We next evaluated mitochondrial activity via the Seahorse MitoStress Test. While LPS and BCR activation elevated basal oxygen consumption rates (a proxy for mitochondrial activity) compared with unstimulated controls, LPS elevated them even more (**Figure 1B**). Overall, LPS reprogrammed B cells toward a highly energetic state, as indicated by elevated OCR and extracellular acidification rate (ECAR, a measure of glycolysis), and αIgM stimulation engendered an intermediate state compared with quiescent, unstimulated B cells (**Figure 1C**). While LPS induced mitochondria to work at maximum capacity, as indicated by a low spare respiratory capacity (SRC), αIgM improved mitochondrial reserve (**Figure 1D**), indicating improved metabolic adaptation. We next interrogated whether changes in mitochondrial respiration were associated with differences in mitochondrial reactive oxygen species (mROS), which could indicate mitochondrial stress. Interestingly, while both activating conditions induced mROS, BCR-dependent stimulation elicited a more pronounced increase (**Figure 1E**), indicating a distinct mitochondrial state compared to LPS. Taken together, these data indicate that TLR4 and BCR signalling engage distinct metabolic programs in B cells.

**Figure 1:**
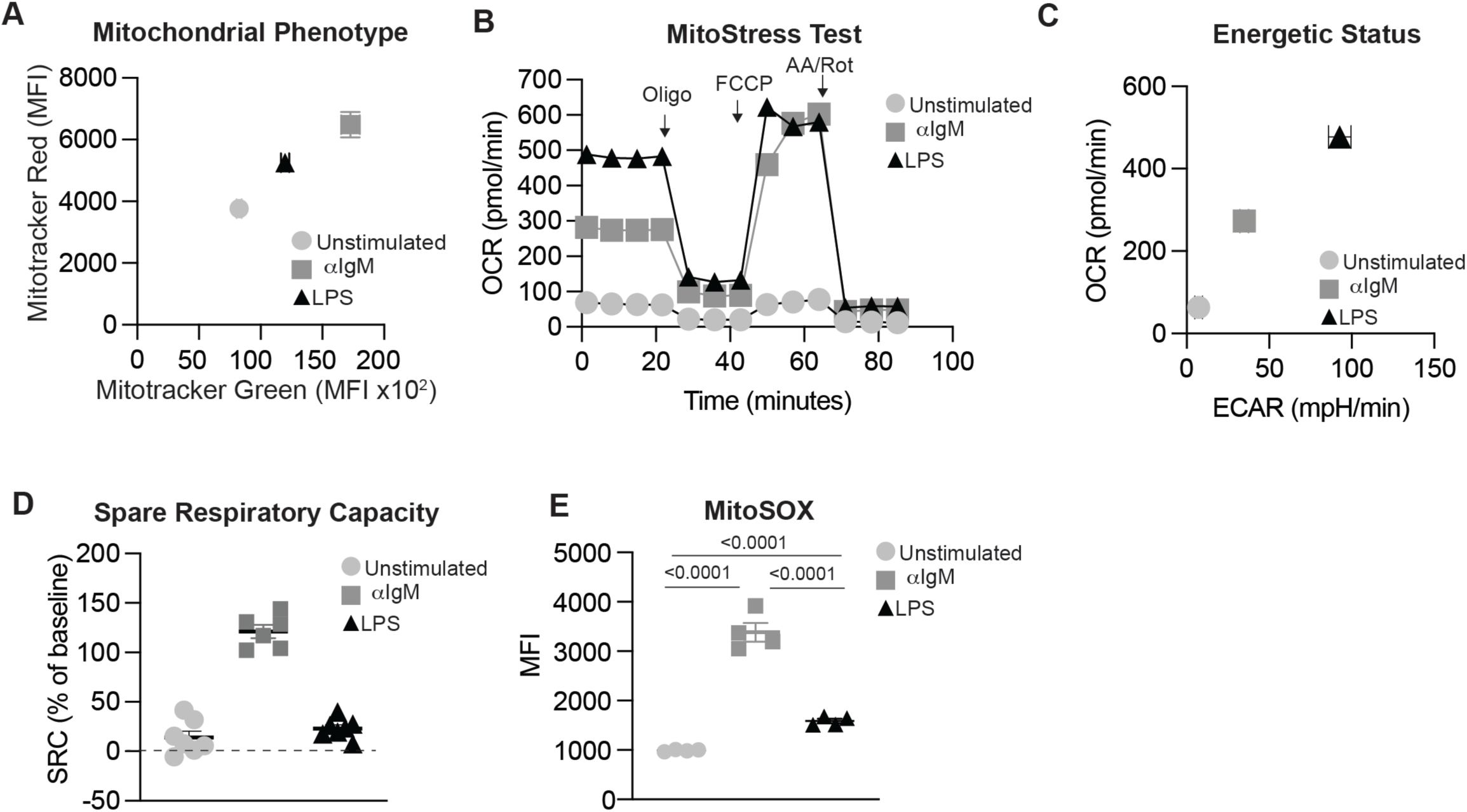
Stimulation-specific mitochondrial remodelling in B cells. Primary B cells were left unstimulated or activated for 24 h with LPS or anti-IgM. **(A)** Mitochondrial mass and membrane potential were assessed by flow cytometry using MitoTracker Green and MitoTracker Red, respectively, and quantified as mean fluorescence intensity (MFI). **(B)** Mitochondrial respiration was assessed using the Seahorse XF Mito Stress Test. Oxygen consumption rate (OCR) was measured at baseline and following sequential injection of oligomycin, FCCP, and rotenone/antimycin A. **(C)** OCR and extracellular acidification rate (ECAR) were compared across unstimulated, LPS-activated, and anti-IgM-activated B cells to assess their metabolic profiles. **(D)** Spare respiratory capacity was calculated as the difference between maximal and basal respiration. **(E)** Mitochondrial reactive oxygen species (mitoROS) were assessed using MitoSOX Red and quantified by flow cytometry as MFI. Data are presented as mean ± SD (n = 3) and are representative of three independent experiments. Statistical significance was determined by one-way ANOVA followed by Tukey’s multiple-comparisons test.

### B cells adapt metabolically to lipid environments and reprogram lipid utilisation upon activation

Given that the microenvironment can shape immune cell metabolism^11,15^, we first evaluated whether B cells responded metabolically to their environment. We first evaluated three immune compartments with descending exposure to adipose tissue (a metabolically active organ): bone marrow, mesenteric lymph nodes (mLN), inguinal lymph nodes (iLN), and the spleen, and assessed lipid droplet levels and expression of the LCFA translocase CD36. Interestingly, in the bone marrow (an environment with the highest direct adipocyte contact), we found that B cells enhanced lipid droplet formation and CD36 expression (**Figure 2A and B**). While lipid droplet accumulation remained unchanged between lymph nodes and the spleen, we detected higher CD36 levels in the mLN than in the iLN and spleen, indicating that B cells adapt to lipid environments (**Figure 2A and B**). We next assessed whether B cells reprogram lipid storage in response to B cell receptor (BCR)-dependent (αIgM and IL-4) and independent (LPS and IL-4) activation. LipidTOX staining showed that BCR stimulation within 24 hours preferentially enhanced lipid storage, as indicated by increased LipidTOX staining compared with LPS and unstimulated B cells (**Figure 2C**). Lipid droplets can be driven by exogenous uptake of saturated fatty acids; therefore, we evaluated BODIPY-conjugated MCFA (lauric acid) and LCFA (palmitate) in unstimulated versus stimulated B cells. Interestingly, while lauric acid uptake was unaffected by activation status, LPS-induced activation increased palmitate uptake compared to BCR engagement (**Figure 2D and E**). Cholesterol metabolism regulates GC B cell responses^16^ and is necessary for lipid droplets. Therefore, we sought to understand if different B cell stimuli differentially modulate cholesterol absorption in B cells. Indeed, both stimuli increased cholesterol acquisition after 24 hours of activation compared with unstimulated controls; however, BCR stimulation robustly enhanced cholesterol uptake compared with LPS, suggesting distinct metabolic programs (**Figure 2F**). Collectively, these data prompted us to assess whether free fatty acid composition responds to activation cues in B cells. Indeed, after 24 hours of activation – either via LPS or BCR engagement – free fatty acid abundance and profile showed a clear shift towards C16 and C18 monounsaturated and polyunsaturated fatty acids, suggesting increased desaturation (**Figure 2G**). Taken together, these data indicate that B cells reprogram their lipid and metabolic profiles in a distinct fashion in response to the type of external stimuli.

**Figure 2:**
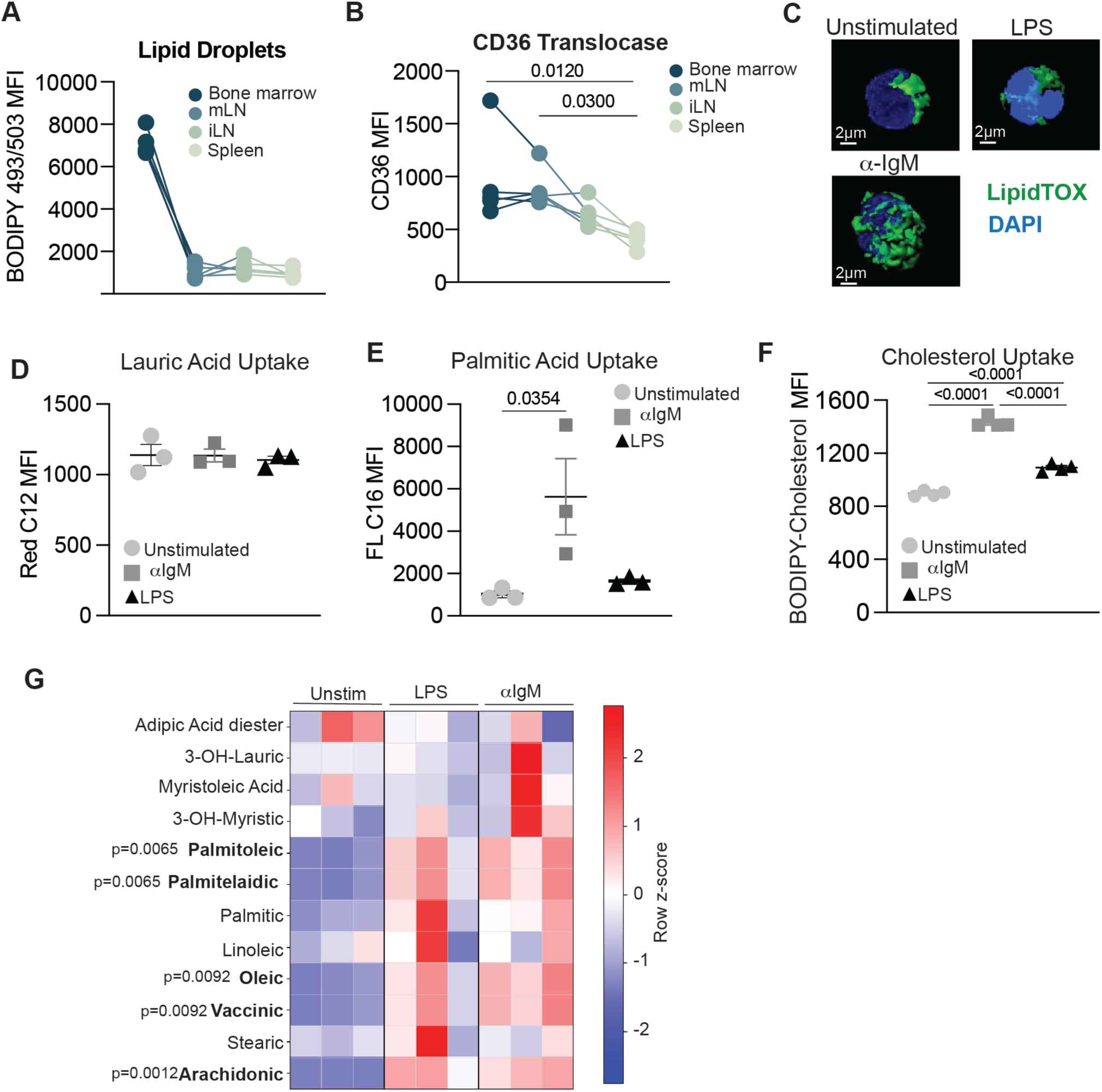
Adipocyte-rich environments reprogram B cells, and upon activation, B cells reprogram their lipid metabolism. **(A and B)** Lipid droplet content **(A)** and CD36 expression **(B)** were assessed by flow cytometry in B cells isolated from the bone marrow (BM), mesenteric lymph nodes (mLN), inguinal lymph nodes (iLN), and spleen of individual male mice (n = 5) and quantified as mean fluorescence intensity (MFI). **(C)** Lipid droplets were visualized by immunofluorescence microscopy in B cells left unstimulated or activated for 24 h with LPS or anti-IgM. Representative Z-stack images are shown. **(D–F)** B cells were left unstimulated or activated for 24 h with LPS or anti-IgM, and lipid uptake was assessed by flow cytometry and quantified as MFI. Uptake of **(D)** lauric acid was assessed using BODIPY Red C12, **(E)** palmitic acid using fluorescent C16 (FLC16), and **(F)** cholesterol using BODIPY-cholesterol. **(G)** Free fatty acids were analyzed by GC-MS, with abundance normalized to the internal standard. For mouse experiments, data are presented as mean ± SEM (n = 5 individual male mice). For in vitro experiments, data are presented as mean ± SD (n = 3) and are representative of three independent experiments. Statistical significance was determined by one-way ANOVA followed by Tukey’s multiple-comparisons test.

### Saturated fatty acid uptake and utilisation is critical for B cell proliferation

In response to immunisation, B cells alter and increase palmitate oxidation^6^. However, lipid metabolism is a complex process of uptake, activation, and utilisation^17^; therefore, we targeted different aspects of saturated-fat metabolism (**Figure 3A**). To test this, we targeted CD36 – part of the LCFA import machinery - with SSO under two stimulatory conditions: BCR-dependent stimulation and LPS stimulation. Dose-dependent inhibition of CD36 profoundly limited B cell proliferation in both BCR- and LPS-activating conditions (**Figure 3B and Figure S1A**). LCFA fatty acids require an acyl-CoA group to be utilised for FAO, membrane incorporation, signalling, or lipid synthesis; the ACSL family regulates this activation^17^. Therefore, we next tested the importance of ACSL function in B cell proliferation with the inhibitor Triacsin C. In a dose-dependent manner, preventing LCFA activation for downstream use significantly abrogated B cell proliferation, indicating that LCFA are essential for B cell expansion (**Figure 3C and Figure S1C)**. Once acyl-CoA has been added to saturated fatty acids, they can then be oxidised in the mitochondria via β-oxidation. Long-chain fatty-acid oxidation requires transport of fatty acyl-CoAs into mitochondria through the CPT-dependent carnitine shuttle. Pharmacological inhibition of CPT1B or CPT1A with oxfenicine or etomoxir, respectively, revealed distinct requirements between activation pathways: LPS-induced lipid metabolic reprogramming depended on both CPT1 isoforms, whereas BCR-dependent activation relied predominantly on CPT1A-dependent fatty-acid oxidation (**Figure 3D-E and Fig. S1C-D**). Together, these data highlight the crucial importance of LCFA uptake, activation and utilisation during B cell activation in both BCR- and TLR-dependent activation.

**Figure 3:**
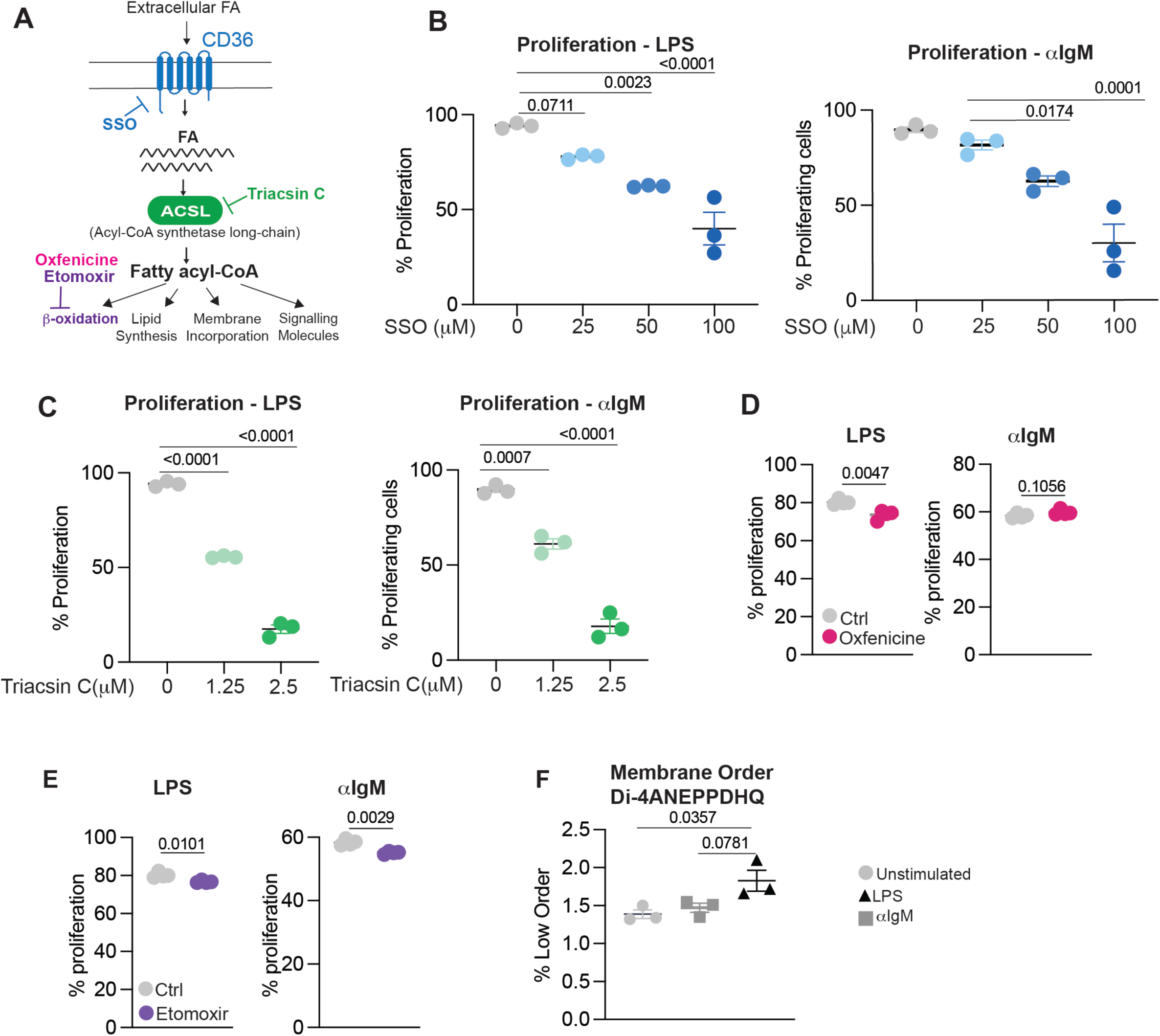
Upon activation, B cells become dependent on lipid uptake, activation and utilisation. B cells were activated with LPS or anti-IgM and proliferation was assessed following pharmacological inhibition of distinct pathways involved in fatty acid uptake and metabolism. **(A)** Schematic representation of the metabolic targets and pharmacological inhibitors used to interrogate fatty acid uptake and metabolism. **(B)** B-cell proliferation following inhibition of CD36 with sulfosuccinimidyl oleate (SSO). **(C)** B-cell proliferation following inhibition of the acyl-CoA synthetase long-chain (ACSL) family with Triacsin C. **(D)** B-cell proliferation following treatment with oxfenicine (500 µM), an inhibitor of CPT1B. **(E)** B-cell proliferation following treatment with etomoxir (5 µM), an inhibitor of CPT1A. **(F)** Plasma membrane lipid order was assessed by flow cytometry using the environment-sensitive membrane probe di-4-ANEPPDHQ, demonstrating decreased membrane order following B-cell activation. Data are presented as mean ± SD (n = 3) and are representative of three independent experiments. Statistical significance was determined by one-way ANOVA followed by Tukey’s multiple-comparisons test.

### Adipocytes and metabolic adaptation to environmental saturated fatty acids

Adipocytes are reliable sources of saturated fatty acids^18^ and are in close proximity to B cells, where this proximity could shape their metabolism (**Figure 2A and B**). However, how adipocytes directly contribute to B cell proliferation and metabolic crosstalk remains unknown (**Figure 4A**). Therefore, we examined the direct effect of adipocytes on B cell proliferation. Using differentiated adipocyte-like 3T3-L1 cells, we performed co-culture assays with primary B cells activated with αIgM and found that adipocytes modestly increased B-cell proliferation (**Figure 4B**). To distinguish direct from indirect effects, we assessed the effect of adipocyte-conditioned medium (ACM) on B-cell proliferation in response to BCR and LPS stimulation. Interestingly, we observed that under both stimuli, in a dose-dependent manner, ACM inhibits B cell proliferation (**Figure 4C and Figure S2A**), suggesting that B cells may provide signals to adipocytes to help with proliferation. Because adipocytes can serve as a source of exogenous fatty acids, we next assessed lauric acid and palmitic acid transfer from adipocytes to B cells and compared it with direct uptake using RedC12 and FLC16, respectively. Strikingly, following BCR stimulation, B cells acquired significantly more lauric acid through adipocyte-mediated transfer than through direct uptake from the extracellular environment (**Figure 4D**). In contrast, palmitate uptake was comparable whether supplied directly or transferred from adipocytes (**Figure 4E**).

**Figure 4:**
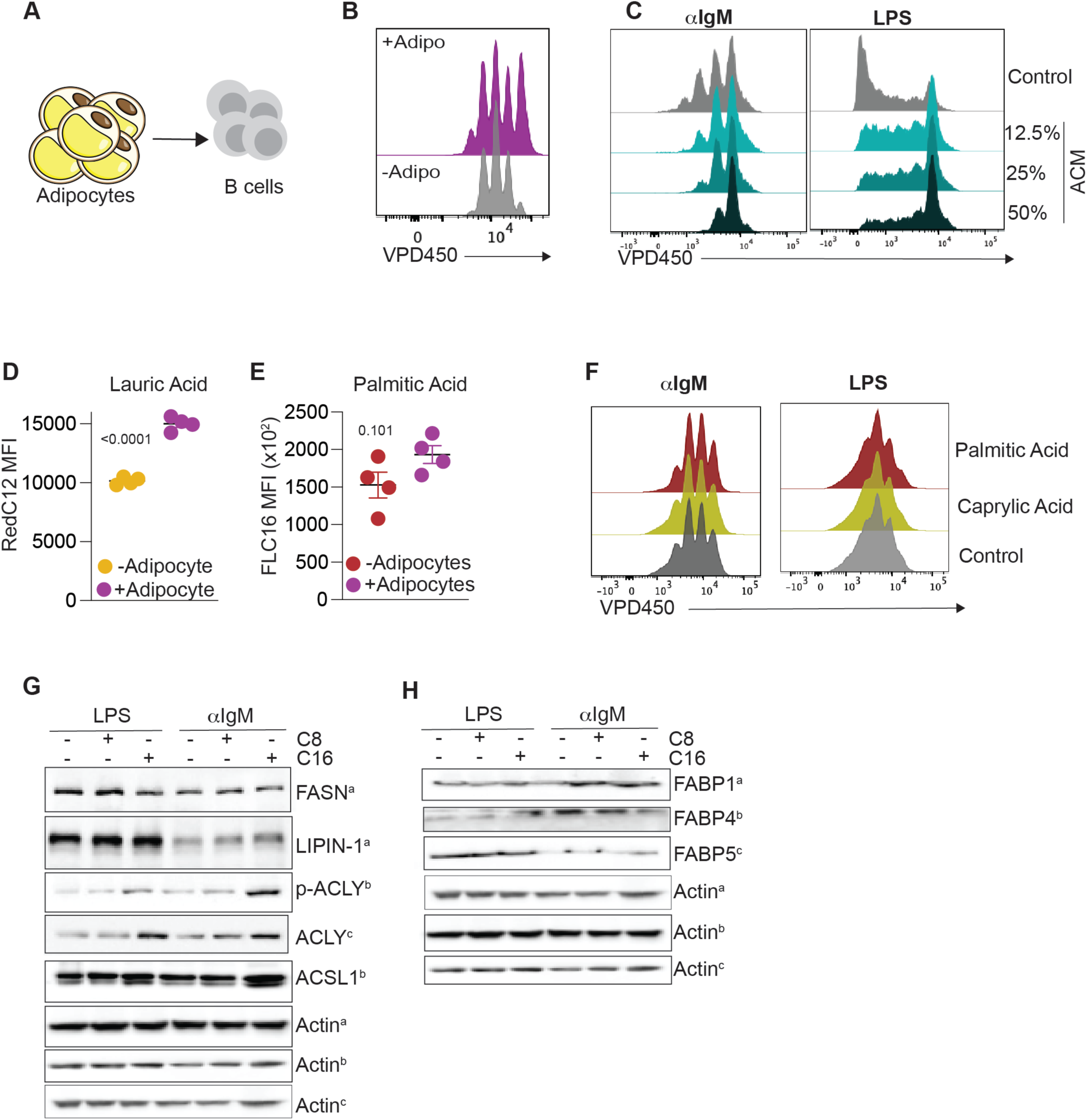
B cells interact with adipocytes and metabolically adapt to exogenous saturated fatty acids. **(A)** Schematic representation of the experimental design used to investigate the effects of adipocytes and adipocyte-derived lipids on B-cell responses. **(B)** Differentiated 3T3-L1 adipocytes were directly co-cultured with B cells activated with anti-IgM for 24 h, followed by assessment of B-cell proliferation on day 3. **(C)** B-cell proliferation following activation in the presence of adipocyte-conditioned medium (ACM). **(D and E)** Transfer of fatty acids from differentiated 3T3-L1 adipocytes to B cells was assessed using adipocytes pre-labeled with fluorescent **(D)** lauric acid (BODIPY Red C12) or **(E)** palmitic acid (fluorescent C16; FLC16). Fatty acid uptake by B cells cultured with or without pre-labeled adipocytes was quantified by flow cytometry. **(F)** B-cell proliferation following activation with LPS or anti-IgM in the presence of caprylic acid (C8:0) or palmitic acid (C16:0). **(G and H)** Western blot analysis of protein lysates from B cells activated with **(G)** LPS or **(H)** anti-IgM in the presence of caprylic acid (C8:0) or palmitic acid (C16:0). Data are presented as mean ± SD. Statistical significance was determined using Welch’s unpaired t-test, as indicated.

We next evaluated whether exogenous MCFA and LCFA modulate B cell proliferation. Culturing BSA conjugated to caprylic acid (MCFA) and palmitate during stimulation revealed no significant effect (**Figure 4F**). Although the readouts are similar between BCR and LPS activation, the pathways supporting metabolic adaptation to saturated fatty acids may differ. Interestingly, ACSL1 expression increased in the presence of palmitate with either BCR or LPS induction, potentially helping redirect LCFA (**Figure 4G**). Surprisingly, we observed increased phosphorylation of ATP-citrate lyase (ACLY) and total levels in activated B cells in the presence of palmitate (**Figure 4G**), suggesting a mechanism to buffer lipotoxic stress^19^. Fatty acid-binding proteins FABP1, FABP4, and FABP5 did not differ in their response to activation in the presence of palmitic or caprylic acid (**Figure 4H**). Together, these findings reveal that B cells differentially acquire and metabolically adapt to exogenous MCFAs and LCFAs, with adipocytes preferentially enhancing MCFA acquisition while LCFAs induce remodelling of lipid-handling pathways without directly altering B-cell proliferation.

### Chain length matters in dietary saturated fatty acids during B cell immunization

Chronic high-fat diet – often referred to as diet-induced obesity – leads to poorer B cell response^10^. However, this does not discriminate between different saturated fat chain lengths, nor does it explain what happens before metabolic dysfunction. To evaluate the effects of individual dietary saturated fatty acids, we generated bespoke diets: control diet (17.4% fat), MCFA-rich diet (where 6% of the 17.4% is replaced with coconut oil), LCFA-rich diet (3% palm oil and 3% cocoa butter replaced of the 17.4%) and a high fat obesogenic diet (60% lard and 35% sucrose) (**Figure 5A**). C57BL/6J mice were fed these four diets for 2 weeks before sheep red blood cell (sRBC) immunisation to induce a GC B cell response in the spleen. At day 8 post-challenge, we assessed GC B cell frequencies in the splenic B cell population and found a significant increase in GC B cells (GL-7^+^CD95^+^) in both the LCFA-rich and obesogenic diets (**Figure 5B**) compared with the control and MCFA-rich diets. Germinal centre B cells undergo dynamic cycling between the dark zone, where they rapidly proliferate and undergo somatic hypermutation, and the light zone, where B cells with higher-affinity B-cell receptors are selected through interactions with follicular dendritic cells and T follicular helper cells. Using CXCR4 (dark zone) and CD86 (light zone) to identify these two zones in the germinal centre, we found that both an LCFA-rich diet and an obesogenic diet increased GC B cells in the light zone (**Figure 5C**). The increased representation of light-zone GC B cells suggests altered B-cell selection within the germinal centre, potentially involving changes in Tfh-mediated help. We next assessed the CD4 Tfh cells in the germinal centre. Strikingly, both the LCFA-rich diet and the obesogenic diet increased CD4 Tfh cell frequencies in the germinal centre (**Figure 5D and Figure S3A**). Collectively, these data suggest that a diet richer in LCFA, independent of obesity, enhances germinal-centre responses characterized by increased GC B cells and Tfh-associated B-cell selection.

**Figure 5:**
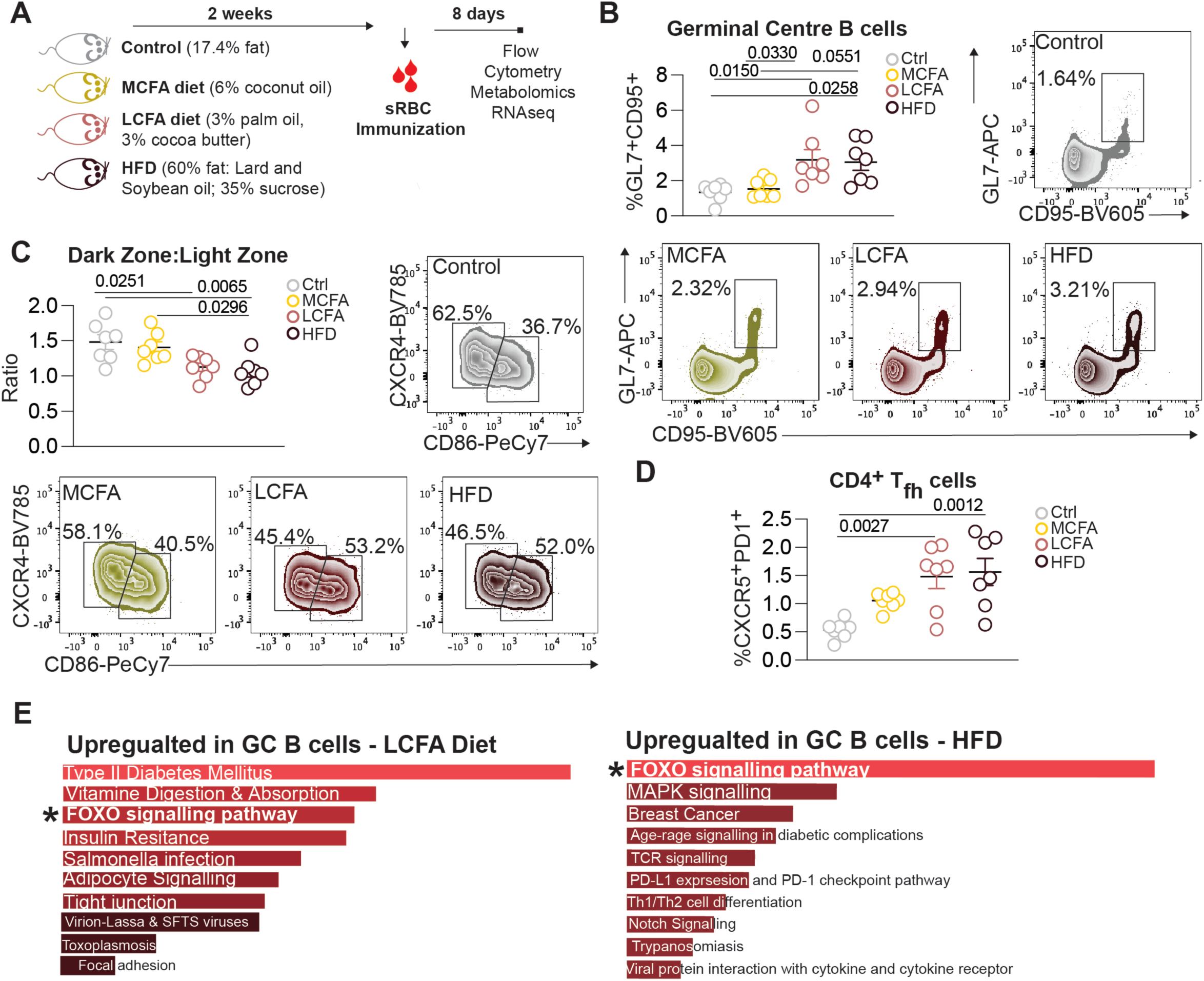
Short-term exposure to diets rich in saturated long-chain fatty acids improves germinal centre B cell responses. **A)** Schematic representation of the experimental design, including dietary intervention and sheep red blood cell (sRBC) immunization timeline. **(B)** Frequency of germinal center (GC) B cells in the spleen at day 8 following sRBC immunization, with representative flow cytometry plots shown. **(C)** Ratio of dark zone (DZ) to light zone (LZ) GC B cells at day 8 post-immunization. Representative flow cytometry plots showing identification of DZ and LZ GC B cells based on CXCR4 and CD86 expression are shown. **(D)** Frequency of T follicular helper (Tfh) cells among splenic CD4+ T cells, identified by co-expression of PD-1 and CXCR5. **(E)** KEGG pathway enrichment analysis of RNA-seq data from splenic GC B cells isolated at day 8 post-immunization. For RNA-seq analysis, n = 3 biological samples per group, with GC B cells from two mice pooled to generate each sample. For **(B–D)**, n = 7–8 mice per group. Data are presented as mean ± SEM. Statistical significance was determined by one-way ANOVA followed by Tukey’s multiple-comparisons test.

Diet may induce metabolic remodelling in GC B cells; therefore, we measured mitochondrial mass and potential in GC B cells generated in mice on the four diets. Unexpectedly, both the LCFA-rich and obesogenic diets reduced the MitoTracker Red/Green ratio, indicating changes in mitochondrial physiology (**Figure S3B**). We further examined diet-induced alterations in GC B cells by performing RNA-seq on GC B cells FACS-sorted from the control, MCFA-rich, LCFA-rich, and obesogenic diets. Interestingly, at the transcriptional level, GC B cells from LCFA-rich and obesogenic diets resemble each other more closely than those from MCFA-rich and control diets (**Figure S3C**). KEGG pathway gene set enrichment analysis of differentially expressed genes identified significant enrichment of pathways associated with FOXO signalling in GC B cells exposed to LCFA-rich and obesogenic diets compared with the control diet (**Figure 5E**). Together, these findings suggest that LCFA-rich and obesogenic diets induce a shared metabolic and transcriptional remodelling of GC B cells, characterized by altered mitochondrial physiology and FOXO-associated signalling

### Dietary fatty acid composition remodels the lipidome and adaptive immune cells

Diet can induce systemic changes in the lipidome and metabolome^20,21^. Performing LC-MS of serum collected at endpoint from all four diets revealed the greatest changes in the lipidome. Relative to control, all three diets induced extensive remodelling of the circulating lipidome (**Figure S4A**). The LCFA-rich diet produced a lipidome more concordant with HFD than with the MCFA-rich diet across several lipid species (**Figure S4B**), with the greatest concordance in triacylglyceride (TAG) species, followed by phosphatidylcholines (PC) (**Figure S4C**). This shared signature was characterised by a significant enrichment of relatively saturated, low-unsaturation TAG species, particularly 18:0 (stearate)- and 18:1(oleate)-containing TAGs, together with coordinated depletion of multiple phospholipid species containing stearic acid (18:0) (**Figure 6A and S4D**). These findings suggest that LCFA and HFD converge on a similar remodelling of lipid metabolism, characterized by enrichment of stearic acid (18:0)-containing neutral lipid species and a concomitant depletion of 18:0-containing phospholipids. This reciprocal pattern is consistent with altered fatty acid partitioning, potentially favouring neutral lipid storage over incorporation into membrane phospholipids.

**Figure 6:**
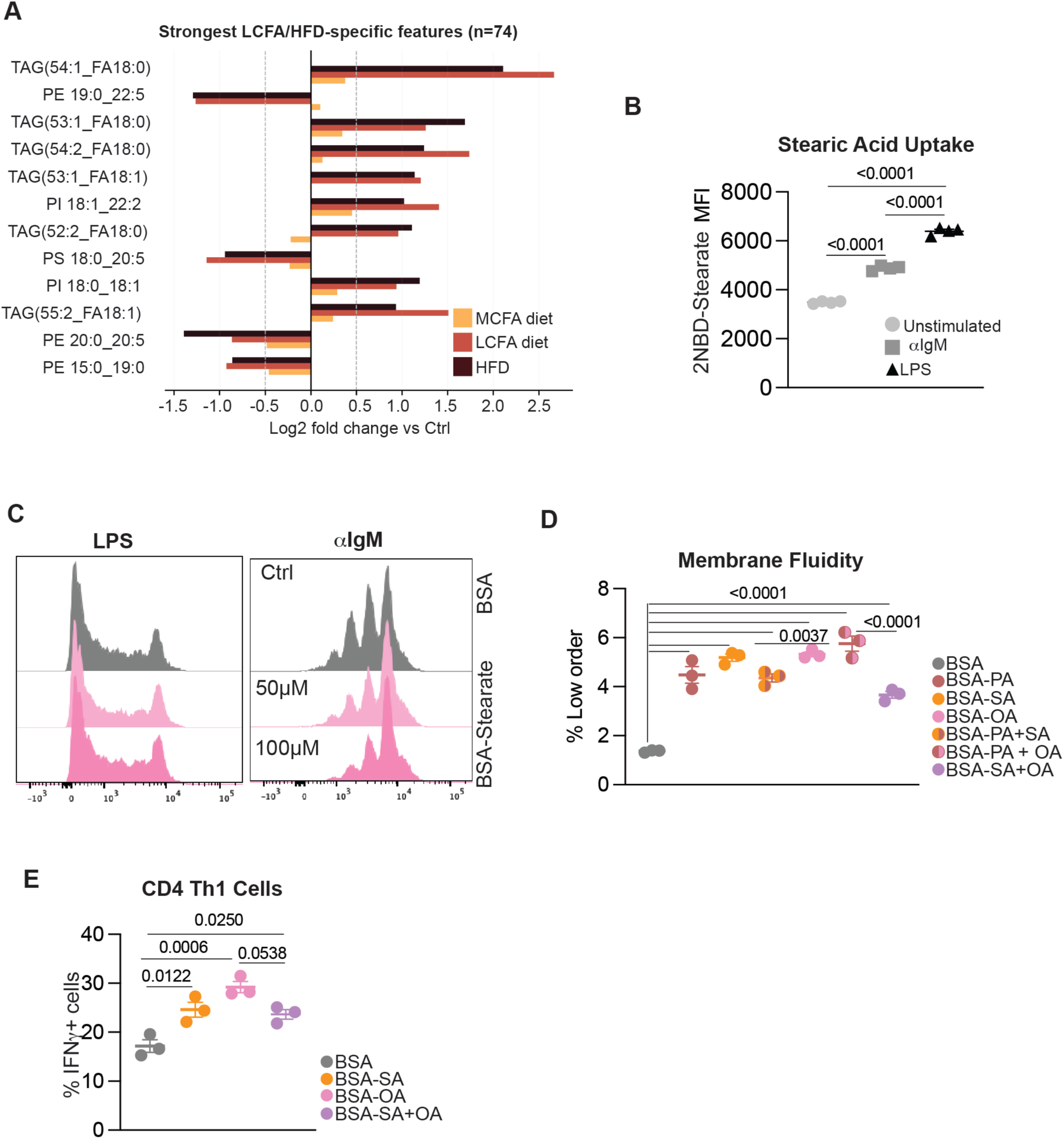
Dietary fatty acids rewire the circulating lipid, and its composition can impact B cell membrane fluidity and T cell differentiation. **(A)** Circulating lipid profiles were analyzed at day 8 following sRBC immunization in mice fed control, medium-chain fatty acid (MCFA), long-chain fatty acid (LCFA), or high-fat (HFD) diets. Lipid species exhibiting the strongest LCFA/HFD-specific changes are shown. Features were selected based on a concordant change in the LCFA and HFD groups relative to control (|log2 fold change| ≥ 0.5), while remaining below this threshold or changing in the opposite direction in the MCFA group (n = 74 lipid features). Bars represent log2 fold changes relative to the control diet for the indicated representative lipid species. **(B)** Stearic acid uptake by unstimulated B cells or B cells activated for 24 h with LPS or anti-IgM was assessed by flow cytometry using fluorescent 2NBD-stearate and quantified as mean fluorescence intensity (MFI). **(C)** B-cell proliferation following stimulation with LPS or anti-IgM in the presence of BSA-conjugated stearic acid (50 or 100 μM). Representative flow cytometry proliferation profiles are shown. **(D)** Plasma membrane lipid order was assessed in anti-IgM-activated B cells cultured in the presence of BSA-conjugated palmitic acid (PA), stearic acid (SA), oleic acid (OA), or the indicated fatty acid combinations. BSA alone served as the vehicle control. Membrane lipid order was assessed by flow cytometry using the environment-sensitive probe di-4-ANEPPDHQ and is shown as the percentage of low-order membrane. **(E)** CD4+ T cells were differentiated under Th1-polarizing conditions for 3 days in the presence of BSA-conjugated SA, OA, or SA+OA, with BSA alone serving as the vehicle control. Th1 differentiation was assessed following restimulation by intracellular cytokine staining for IFN-γ, and data are shown as the frequency of IFN-γ+ CD4+ T cells. For **(A)**, lipid features quantified in at least 4 of 8 mice per dietary group were retained for analysis, and group-level log2 fold changes were calculated relative to the control diet. For **(B–E)**, data are presented as mean ± SD and statistical significance was determined by one-way ANOVA followed by Tukey’s multiple-comparisons test.

Fatty acids are stored as neutral TAGs, allowing for future use while avoiding the consequences of lipid toxicity^22^. Given the enrichment of stearate in response to LCFA and HFD during immunisation, we sought to test the direct effect of stearate on B cells. Both LPS- and anti-IgM-stimulated B cells exhibited increased stearate uptake compared with unstimulated cells, with the greatest uptake observed following LPS stimulation (**Figure 6B**). Given this enhanced uptake, we next asked whether stearate availability directly influenced B-cell proliferation. Despite its increased acquisition by activated B cells, exogenous stearate did not significantly alter proliferation under either stimulatory condition (**Figure 6C**).

Serum lipidomics suggest that changes in lipid partitioning induce a converging phenotype in the LCFA-rich and obesogenic diet, where 18:0-containing phospholipids are depleted, suggesting they are diverted away from structural membrane lipids (**Figure 6A**). These changes could directly affect membrane fluidity in B cells. Membrane fluidity and order are biophysical properties of the plasma membrane determined in part by lipid composition, with changes in lipid packing influencing membrane organisation, receptor signalling, and cellular function^23^. Membrane fluidity is essential for BCR-dependent signal transduction in B cells^24,25^. Therefore, we tested how exogenous palmitate, stearic acid, oleate, and their combinations directly affect membrane order in BCR-stimulated B cells. Interestingly, all exogenous saturated fatty acids and MUFAs or in combination, increased low-order membranes – an indicator of less-packed, more fluid membranes. Taken together, these data indicate that exogenous fatty acids may affect the membrane structure of activated B cells.

Finally, sRBC immunisation revealed that lipid changes could also affect CD4 T cells, either indirectly through improved GC B cells or directly. We tested how LCFA and the MUFA oleate directly affected CD4 T helper 1 cells. Interestingly, oleate along had a profoundly promoted differentiation, as well as stearic acid alone. However, the combination, while significantly compared to BSA controls, was modestly less than oleic alone, suggesting that lipid handling during CD4 T cell differentiation may impact their polarisation. Taken together, diet-induced changes in the lipidome have far-reaching effects on immune cells in response to an external stimulus.

## Discussion

Our findings identify fatty-acid availability and chain length as determinants of B-cell metabolic adaptation and demonstrate that dietary lipid composition can shape germinal-centre responses independently of established obesity. Here, we show that fatty-acid metabolism is dynamically regulated during B-cell activation and differs by activating signal and extracellular lipid environment. BCR- and TLR4-dependent activation induced distinct mitochondrial and lipid-handling programs, while disruption of LCFA uptake, activation, or mitochondrial utilisation impaired B-cell proliferation. While LPS-induced activation of B cells engages in glycolysis to promote lipogenesis^26^, our data also underpin an important role in lipid uptake, which could converge on fatty acid utilization mediated by ACSLs for fatty acid oxidation.

B-cell activation does not simply depend on lipid acquisition globally but instead induces stimulus-dependent changes in the uptake, storage and composition of cellular lipids. Interestingly, we and others have shown that either BCR or TLR activation of B cells enhances intracellular MUFAs^27^, such as oleic acid. We also observed TAGs enriched in oleic acid in LCFA and high-fat diets, where GC B cell frequencies improved. Given that stearoyl-CoA desaturase is essential for B cell responses^27^, B cells may shuttle saturated fatty acids towards oleic acid production for expansion and function. Therefore, while GC B cells depend on fatty acid oxidation and rely on the sterol regulatory binding protein (SREBP) pathway to mediate robust B cell responses^6,28^, fatty acid partitioning towards MUFAs is also essential for B cell expansion.

One way modifying lipid partitioning can affect function is through membrane fluidity. Increased membrane fluidity – when plasma membranes are less packed due to increased MUFA and PUFA – will improve receptor signalling. Therefore, this in vivo and in vitro increase in MUFAs could reflect cells repartitioning lipids for membrane composition. Indeed, we found that exogenous LCFA, MUFA or the combination promotes membrane fluidity within 24 hours of BCR engagement. Another route for lipid handling is fatty oxidation, and we found that overall inhibition of either CPT1A or CPT1B using etomoxir and oxfinecine, respectively, limited B cell proliferation post-stimulation. Interestingly, loss of CPT2 does not influence B cell development, GC B cell response or antibody production^29^. However, excess exogenous fatty acids, as seen in diet-induced obesity, increase fatty acid oxidation in CD4 T cells^30^, and therefore, modulating this pathway via diet for B cell responses could also evoke a similar response. Indeed, our data indicate that short-term exposure to changes in dietary saturated fat is sufficient to improve GC B cell frequencies. However, whether this occurs through increased fatty acid oxidation remains to be determined. However, obesity was found to induce metabolic stress in CD4 T cells and elevate PI3K signalling, favouring effector memory cell fate^31^. Indeed, RNA-seq data from sorted GC B cells from immunised mice on different diets indicated enrichment in FOXO signalling in both the LCFA and high-fat diets compared with the control diet. Since FOXO is a target of AKT, and PI3K is upstream of this pathway, this mechanism could also be in play in B cells responding to changes in dietary saturated fats.

However, timing could play a salient role since obesity is associated with poor B cell development and antibody responses^10^. We have shown that either altering the composition of saturated fatty acids or increasing caloric intake from fats for a couple of weeks was sufficient to improve GC B cell frequencies after sRBC immunisation. These differences also improved CD4 Tfh frequencies in the spleen, overall suggesting improved responses. Because lipidomics showed that the LCFA and high-fat diet converge on increased TAGs with stearate and oleic acid (MUFA), we directly tested whether these fatty acids affected CD4 T cell differentiation. Indeed, these fatty acids, either alone or combined, improved IFNg production, with oleic acid alone evoking the strongest response. Overall, these findings indicate that the timing and composition of fat matter for improving GC B cell frequencies, as well as CD4 T cells, and could be leveraged during immunisation.

Adipocytes can also serve as a source of exogenous fatty acids. It has been well characterized that B cells promote the adipocyte-obesity-metabolic dysfuction axis^12,32,33^, yet whether adipocytes directly affect B cells outside of the context of obesity was poorly understood. Here we show that adipocytes can metabolically interact with B cells by directly transferring saturated fatty acids to B cells during activation. In addition, we show that this interaction is favourable for B cell expansion, yet adipocyte-conditioned media – bypassing direct interactions-inhibits B cell proliferation. Therefore, it is possible that B cells may signal to adipocytes for metabolic support, given that memory CD8 T cells utilize the adipose tissue as a reservoir and metabolically adapt to lipid-loaded adipocytes^11^.

Collectively, our findings demonstrate that B cells dynamically regulate fatty-acid uptake, activation, utilization and partitioning in response to both activation signals and extracellular lipid availability. They further reveal that dietary fatty-acid chain length can remodel the circulating lipid environment and GC response, with LCFA-rich feeding reproducing several features observed following obesogenic feeding. These findings broaden the role of fatty acids in B-cell biology beyond their function as metabolic fuels and highlight the importance of lipid composition and metabolic fate in shaping B-cell responses.

### Study Limitations

While this study advances our understanding of lipid handling in B cells and how dietary fats can reshape immunisation. However, the exact molecular mechanism driving improved responses with diets richer in saturated LCFA remains unclear. The circulating lipid profile suggests a potential difference in lipid partitioning; this study does not address whether this occurs at the cellular level in B cells. Ex vivo lipidomics of GC B cells isolated from mice immunised and exposed to different dietary fats would help address this question. In addition, this study is limited in understanding the precise window in which diets rich in long-chain saturated fats remain beneficial rather than detrimental.

## Methods

### Isolation and activation of Primary lymphocytes

Primary naïve B lymphocytes were isolated from the spleens of C57BL/6J mice. Spleens were mechanically dissociated through a 70-μm cell strainer in PBS, and splenocytes were counted using methylene blue. B cells were isolated by immunomagnetic negative selection using a commercial isolation kit (19854, STEMCELL Technologies) according to the manufacturer’s instructions. B cells were activated for 24 h with IL-4 (10 ng/mL) in combination with either LPS (5 μg/mL) or anti-IgM (10 μg/mL) in complete RPMI medium containing 10% FBS (Wisent Bio Products), 1% penicillin-streptomycin (10,000 U/mL; Wisent Bio Products), and 0.1% 2-mercaptoethanol (55 mM; Gibco).

Primary CD4□ T cells were isolated from the spleens and lymph nodes of C57BL/6 mice. Single-cell suspensions were prepared and CD4□ T cells were isolated using a CD4□ T-cell isolation kit (19852, STEMCELL Technologies) according to the manufacturer’s instructions. T cells were activated on plates pre-coated with anti-CD3 (clone 145-2C11; BioLegend, 100312) and anti-CD28 (clone 37.51; BioLegend, 102112).

### Differentiation of 3T3-L1 pre-adipocytes

The 3T3-L1 murine preadipocyte cell line was maintained in complete DMEM (Wisent Multicell) supplemented with 10% fetal bovine serum (FBS) and 1% penicillin/streptomycin (P/S). For adipocyte differentiation, cells were seeded in 24-well plates and cultured until approximately 80% confluent. Differentiation was initiated by replacing the maintenance medium with complete DMEM supplemented with 0.5 mM 3-isobutyl-1-methylxanthine (IBMX; Acros Organics), 0.5 µM dexamethasone (Gibco), 10 µg/mL insulin (Sigma-Aldrich), and 20 µM troglitazone (Sigma-Aldrich). On days 4, 6, and 8, the differentiation medium was replaced with complete DMEM supplemented with 10 µg/mL insulin. On day 10, the medium was replaced with standard complete DMEM and cells were maintained thereafter.

Unless otherwise indicated, cells were incubated at 37°C in an incubator humidified with 5% CO□.

### Proliferation assays

Spleens were harvested from C57BL/6 mice and mashed through a 70 µm filter. The splenocytes were resuspended in 1X PBS and counted using methylene blue. Primary B cells were isolated by negative selection using the magnetic separation kits from Stemcell^TM^ according to the manufacturer’s protocol. The isolated primary B cells were washed with PBS and incubated with 1 µM VPD for 10 minutes at 37□ in the dark. The cells were resuspended in complete RPMI media (containing 10% heat-inactivated fetal bovine serum, 1% penicillin-streptomycin, and 0.1% 2-mercaptoethanol). Cells were seeded in 96-well plates at a density of 400 000 cells in 200 µL(2×10^6^ with IL-4(20ng/ml) and LPS(5µg/ml) or □-IgM (10µg/ml) and incubated for 72 hours. Cells were treated with the following exogenous fatty acids: BSA–caprylic acid (C8; Cayman Chemical), BSA–palmitate (C16; Cayman Chemical), and BSA–stearate (C18; Cayman Chemical) at final concentrations of 50 µM and 100 µM. Cells were also treated with Sulfo-N-succinimidyl oleate (Cayman Chemical), triacsin C (Santa Cruz), adipocyte conditioned media, etomoxir (5µM), and oxfenicine (0.5mM).After 72 hours, cells were stained with Viability marker (APC-eFluor 780) for 10 minutes at 4□, then incubated with CD45R, CD19, CD40, CD86, and CD44 for 30 minutes at 4□, then washed and resuspended in 100 µL of FACS buffer for flow cytometry analysis.

### Primary B cells co-cultured with adipocytes

Primary B cells were isolated from the spleen of C57BL/6 mice and activated for 24 hours. The cells were then plated at a density of 2×10^6^ cells /ml in of RPMI complete medium under the following conditions: (i) IL-4(20 ng/ml) and (ii) IL-4(20 ng/ml) + □-IgM (5µg/ml). 3T3-L1 adipocytes were incubated at 37□ overnight with (i) 20µM Lauric acid, (ii) 5µM Palmitate. Both cell types were washed and B cells resuspended in complete RPMI were added to the 3T3 L1 wells and incubated at 37□. After 24 hours, The B cells were washed and stained with a viability marker (APC-eFluor 780) for 10 mins at room temperature. Cells were washed in FACS buffer and resuspended in buffer containing CD19(1/500), CD40(1/800), CD86(1/500), and CD44(1/500). Cells were incubated for 30 minutes at 4□, then washed and resuspended in 100 µL of FACS buffer for flow cytometry analysis.

### Seahorse assays

Primary B lymphocytes were incubated for 24 hours in complete RPMI with and without activators IL-4 and LPS or IL-4 and anti-IgM. The cells were then transferred to Seahorse XF DMEM media(103575-100, Agilent Technologies) containing 12 mM glucose, 2mM sodium pyruvate, and 2 mM glutamine. Cells were seeded at 500 000 cells/well in 96 wells precoated with Cell-Tak(CLS354240, Millipore Sigma), then incubated at 37 °C without CO□ for 1 hour. The mitochondrial modulators (Etomoxir, oligomycin, FCCP, and antimycin A) were prepared to obtain a final concentration of 2.5 µM per well.

## Flow cytometry

### Viability staining

The cells were incubated for 10 minutes at 4°C with 50 µL of eFluor 780 viability dye diluted 1:1000 in 1X PBS. The cells were then washed with 1X PBS and centrifuged at 1300 rpm for 5 minutes.

### Mitochondrial mass and potential

Cells were seeded and incubated for 24 hours in complete RPMI medium. Cells were harvested by centrifugation (1300 rpm, 5 minutes), and viability staining was done as described above. The cells were then incubated for 30 minutes at 37°C (5% CO_2_) in the dark with MitoTracker Green (20 nM) and MitoTracker Deep Red (50 nM) (Invitrogen), diluted in 1X PBS. After washing, the cells were resuspended in their growth medium and stored at 4°C until acquisition.

### Absorption of Fatty acids

Primary B cells were activated with IL-4 and anti-IgM and incubated for 24 hours with and without BODIPY-conjugated lauric acid (BODIPY 558/568 C12, Invitrogen), palmitate (BODIPY FL C16, Invitrogen), stearate (44253, Cayman Chemicals), and cholesterol (GC42964, Good Laboratory Practise Bioscience). The cells were washed and resuspended in PBS with respective BODIPY dyes at a final concentration of 1uM and incubated for 1-2 hours at 37°C. Cells were incubated with viability dye for 10 minutes in the dark, washed and incubated for 30 minutes at 4°C with other external antibodies.

### Lipid droplets and lipid peroxidation

Primary B cells were activated and seeded as described above and incubated for 24 h in standard medium or medium supplemented with BSA-conjugated caprylic acid or palmitate (50 μM). The medium was then replaced with fresh medium containing either the lipid droplet marker BODIPY 493/503 (0.5 μM) or the lipid peroxidation marker BODIPY 581/591 C11 (0.5 μM; Invitrogen), and cells were incubated for an additional 2 h.

Bone marrow cells were isolated from the tibiae and femora of C57BL/6 mice (8–10 weeks old) by flushing the bones with cold HBSS using a 25-gauge needle. Cells were resuspended in PBS and adjusted to 1 × 10□ cells/mL. Mesenteric lymph nodes (mLNs) and inguinal lymph nodes (iLNs) were collected and mechanically dissociated through a 70-μm cell strainer in cold PBS. Cells isolated from the bone marrow, lymph nodes, and spleen were incubated with BODIPY 493/503 (0.5 μM) for 2 h, then stained with an APC-conjugated anti-CD36 antibody at 4°C for 30 min.

### LipidTOX in primary B cells

Cells were fixed with 4% paraformaldehyde (PFA) for 10 min. Following fixation, cells were permeabilized with 0.1% saponin for 30 min at room temperature (RT). Cells were then stained with LipidTOX (Thermo Fisher Scientific, #1043673; 1:200) for 1 h at RT. After washing, nuclei were stained with DAPI (Bioshop Canada, #DAP444.5; 1:10,000) for 10 min. Following a final wash, coverslips were mounted using ProLong Diamond Antifade Mountant (Thermo Fisher Scientific, #P36965). Images were acquired using a Leica Stellaris STED inverted confocal super-resolution microscope and analyzed using Imaris software (Bitplane, version 9.1.2).

### Membrane order (Di-4-ANEPPDHQ assay)

Activated primary B cells were incubated with a fixable viability dye (BD Fixable Viability Stain eFluor 780; 1:1,000 dilution in PBS) and anti-B220-BV605 antibody (1:500 dilution) in a final volume of 50 μL per well. Following surface staining, cells were washed with PBS then stained with 2 μM Di-4-ANEPPDHQ diluted in RPMI or PBS and incubated at 37°C for 30 min. Following staining, cells were incubated for an additional 15 min at room temperature. Cells were analyzed immediately by flow cytometry without an additional wash or fixation step. Di-4-ANEPPDHQ fluorescence was detected using the appropriate fluorescence channels for its green (∼570 nm) and red (∼630 nm) emission signals.

### Th1 cell differentiation

CD4 ^+^ T Cells were cultured for 6 days in T-cell medium (IMDM supplemented with 10% FBS, 1% P/S, and 50µM β-mercaptoethanol). On day 0, culture plates were coated with 5 µg/mL of anti-CD3 (Biolegend, 145-2C11, 1015436) and 2 µg/mL anti-CD28 (Biolegend, 37.51, 14-0281-85) overnight at 4°C or for 2 h at 37°C. The isolated CD4 T cells were then seeded at 1 × 10^6^ cells/well in the coated plates with 10 ng/mL IL-2 and cultured for 3 days, followed by incubation with PMA (1ng/ml), ionomycin (10µg/ml), and GolgiStop for 4 h at 37°C and 5% CO□, followed by intracellular cytokine staining for IFNγ and TNF.

### Western Blot

Primary B cells were seeded at 2 × 10^6^ cells/ml in 6 cm dishes (5 mL total volume) and activated for 24 hours with IL-4 (20 ng/ml) and LPS (5 µg/ml) or anti-IgM (10 ug/ml). Protein lysates were prepared in RIPA buffer (Millipore Sigma) supplemented with a protease/phosphatase inhibitor cocktail (Cell Signaling; 1:100), 10% SDS (1:100), and 1 M benzamidine hydrochloride (1:2,000; Millipore Sigma). The lysates were vortexed and then mechanically sheared using an insulin syringe (BD). After centrifugation, the protein concentration was determined by the Bradford assay.

For the analysis, 40 µg of protein per sample was separated on a 10% SDS-PAGE gel and then transferred to nitrocellulose membranes by wet transfer. After blocking, the membranes were incubated with primary antibodies against fatty acid and lipid metabolism proteins, as well as with an anti-β-actin antibody used as a loading control. After incubation with HRP-conjugated secondary antibodies, the signals were detected by chemiluminescence using the West Femto substrate and acquired on an Azure c6000 imaging system. Membranes were stripped (cat # 91925, Cell Signaling) according to the manufacturer’s protocol.

### In vivo sRBC immunization

C57BL/6J mice (8–10 weeks old) were fed one of four diets for 2 weeks: control (17.4% fat), medium-chain fatty acid (MCFA; 6% coconut oil), long-chain fatty acid (LCFA; 3% palm oil and 3% cocoa butter), or high-fat diet (HFD; 60% fat and 35% sucrose). Mice were immunized intraperitoneally with 2 × 10□ sheep red blood cells (sRBCs) prepared in HBSS and administered in PBS. Spleens were collected 8 days post-immunization, and germinal center (GC) B cells were isolated. GC B cells were stained with antibodies against B220, GL-7 (clone GL7), and CD95 (clone SA367H8) for flow cytometric analysis. Samples were acquired using a BD LSRFortessa flow cytometer and analyzed using FlowJo v10.10.1. Isolated GC B cells were additionally used for metabolomic and RNA-sequencing analyses.

### RNA sequencing

Poly(A) mRNA enrichment, fragmentation, and cDNA library preparation was done from total RNA, followed by adapter ligation, amplification, and purification. Libraries were quality-controlled and sequenced on an Illumina platform. Raw reads were quality-filtered using fastp and mapped to the reference genome using HISAT2 (v2.2.1). Read counts for each gene were generated using featureCounts (v2.0.6), and differential expression analysis was performed using DESeq2 (v1.42.0) for biological replicates. Differentially expressed genes were identified using an adjusted *P* value ≤ 0.05 and |log2 fold change| ≥ 1. GO and KEGG pathway enrichment analyses were performed using clusterProfiler (v4.8.1).

### Free fatty acid metabolomics

Primary B cells were activated for 24 h as described above, and equal numbers of control, LPS-stimulated, and IgM-stimulated cells (10 × 10□ cells per sample) were collected and washed with 0.9% NaCl. Cell pellets were extracted with 80% methanol and dichloromethane, with d27-myristic acid added as an internal standard. The organic fraction was dried, resuspended in pyridine, and derivatized with MTBSTFA containing 1% TBDMCS at 70°C for 60 min. Derivatized samples were analyzed by GC-MS using an Agilent Intuvo 9000 GC coupled to a 5977B mass spectrometer. Fatty acids were identified and quantified based on retention times and diagnostic ions from authentic standards. Raw data were processed using ProteoWizard and the xcms package in R (v4.4.3), and fatty acid peak areas were normalized to the d27-myristic acid internal standard.

### Serum lipidomic profiling

Serum samples collected from mice after dietary treatment were submitted to Linearis Labs (Canada) for targeted metabolomic and lipidomic profiling using the GigaKit™ Metabolomic Panel. The assay quantified up to 1,780 metabolites across 38 biochemical classes using liquid chromatography-tandem mass spectrometry (LC-MS/MS) and direct flow injection-tandem mass spectrometry (DFI-MS/MS). Briefly, serum samples underwent pre-column derivatisation with phenylisothiocyanate (PITC) for amine-containing metabolites and 3-nitrophenylhydrazine (3-NPH) for organic acids and selected lipids. Derivatised samples were analysed in multiple reaction monitoring (MRM) mode on a Waters UPLC Acquity I-Class system coupled to a Xevo TQ-Absolute mass spectrometer, with five analytical runs per sample. Data acquisition, processing, and quantification were performed using Waters Connect software. Metabolites analysed by LC-MS/MS were quantified using seven-point calibration curves based on analyte-to-internal standard (ISTD) peak intensity ratios, whereas lipids and acylcarnitines were semi-quantitatively assessed by DFI-MS using single-point calibration with representative analyte and ISTD standards for each chemical class. Quality control pooled serum samples were included in each analytical batch to monitor measurement variability and facilitate data normalisation. The analytical laboratory assessed data quality using an automated quality control pipeline.

## Acknowledgements

We acknowledge the Animal Facility and Flow Cytometry Facility at the Centre de recherche de l’Hôpital Maisonneuve-Rosemont (CR-HMR) for their technical expertise and support. This work was supported by the Cancer Research Society Operating Grant (Award ID 1057230) and the Cole Foundation. J.B. is supported by a Fonds de recherche du Québec (FRQ) Junior 1 salary award; E.B.G. by the Stavros Niarchos Foundation; B.N. by an FRQ Master’s studentship; and L.B. by a Cole Foundation postdoctoral fellowship.

## Author Contributions

J.B. and F.Z. conceived and designed the study, contributed to experimental work and data interpretation, and J.B. wrote the manuscript. B.N., L.B., E.B.G., and I.G. performed in vtor experiments and analysed data. L.B. and A.L.C. performed the in vivo immunisation experiments with assistance from M.L., E.B.G., and J.B. T.K.N. performed western blots. M.L. performed immunofluorescence and analysis. Y.B. performed and analysed membrane fluidity experiments. C.V. performed metabolomics and analysis. All authors reviewed and approved the final version of the manuscript and agreed to its submission.

