## supplemental file for "Fatty acid metabolism and dietary lipid composition shape B-cell responses"

**Supplementary Figures**

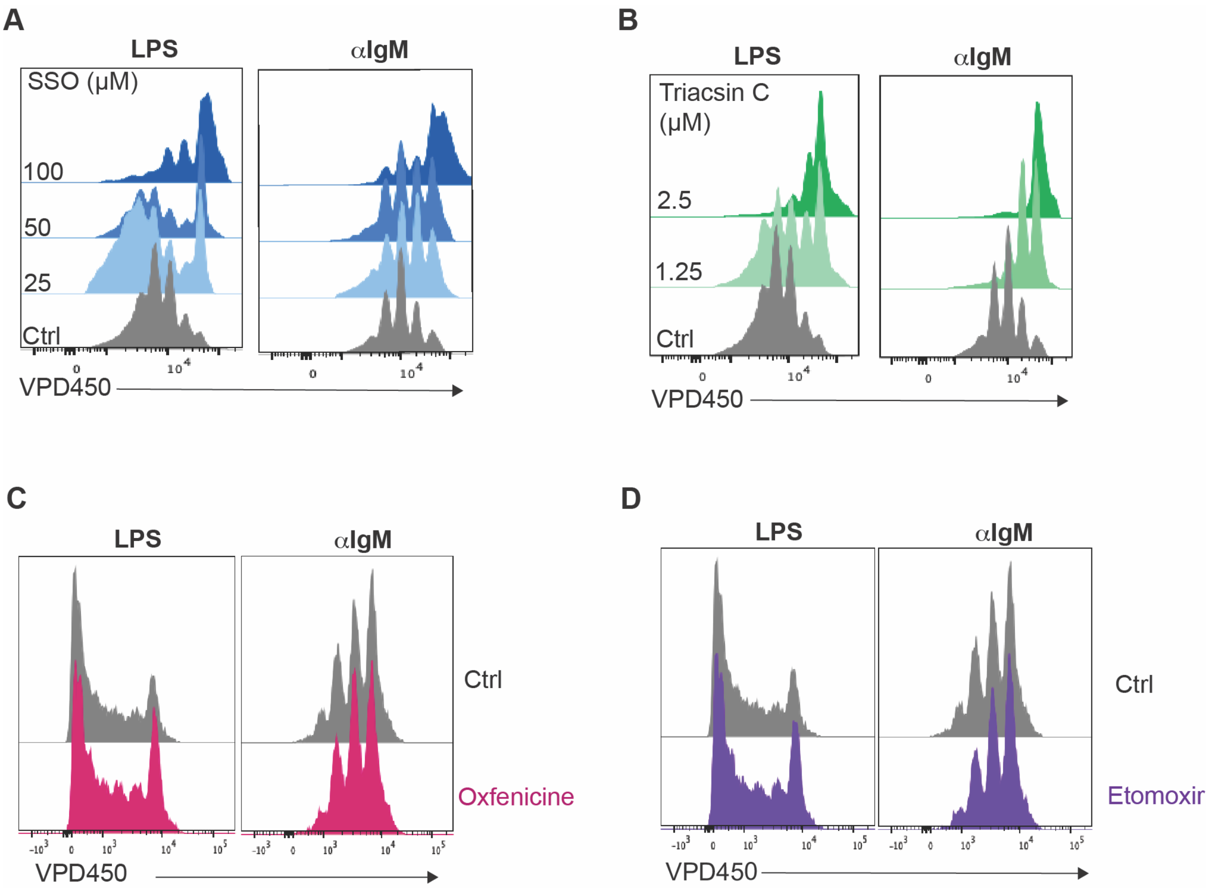


**Figure S1: Target fatty acid metabolism impacts B cell proliferation**

B cells were stimulated with LPS or anti-IgM, and proliferation was assessed by Violet Proliferation Dye dilution and **analyzed by flow cytometry**. **(A)** B-cell proliferation following inhibition of CD36 with sulfosuccinimidyl oleate (SSO). **(B)** B-cell proliferation following inhibition of the acyl-CoA synthetase long-chain (ACSL) family with Triacsin C. **(C)**B-cell proliferation following treatment with the CPT1B inhibitor oxfenicine (500 µM). **(D)** B-cell proliferation following treatment with the CPT1A inhibitor etomoxir (5 µM). Representative Violet Proliferation Dye dilution profiles are shown for LPS- and anti-IgM-stimulated B cells.

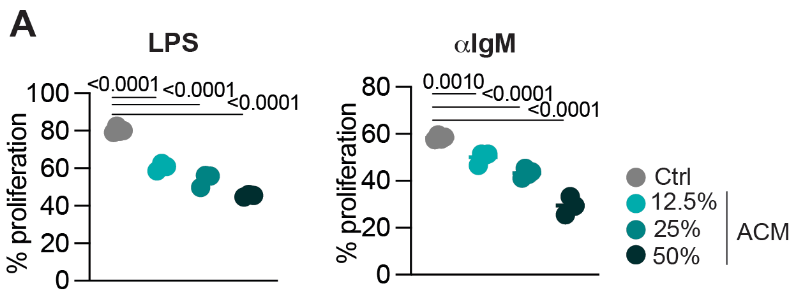


**Figure S2: Adipocyte conditioned medium limits B cell proliferation**

B cells were stimulated with LPS or anti-IgM in the presence of decreasing concentrations (% v/v) of adipocyte-conditioned medium (ACM) collected from differentiated 3T3-L1 adipocytes. B-cell proliferation was assessed by Violet Proliferation Dye dilution and analyzed by flow cytometry. Data are presented as the frequency of proliferated B cells (mean ± SD) and are representative of three independent experiments. We determined statistical significance by one-way ANOVA followed by Dunnett's multiple-comparisons test.

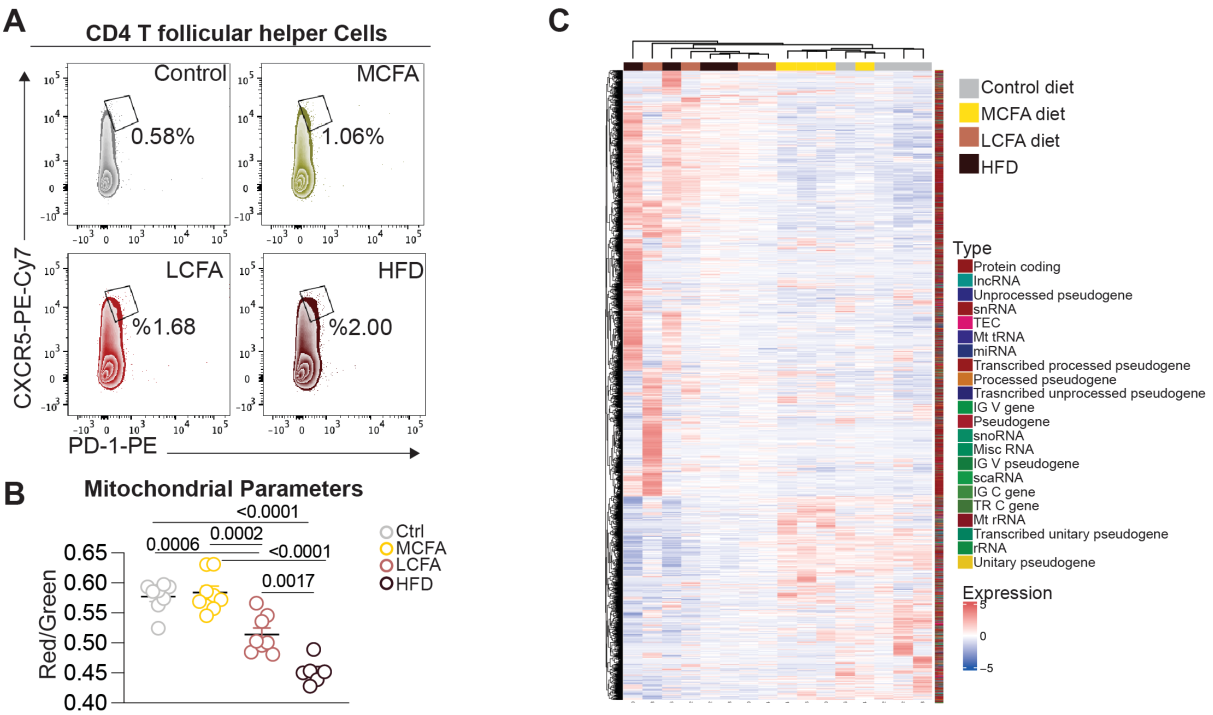


**Figure S3: Diets rich in long-chain fatty acids promote CD4 T follicular helper cells and reprogram the germinal centre B cell transcriptome.**

Mice were maintained on control, medium-chain fatty acid (MCFA), long-chain fatty acid (LCFA), or high-fat (HFD) diets and immunized with sheep red blood cells (sRBCs). Splenic immune responses were assessed at day 8 post-immunization. **(A)** Representative flow cytometry plots showing T follicular helper (Tfh) cells within the splenic CD4+ T-cell population, identified by co-expression of PD-1 and CXCR5. **(B)** Mitochondrial parameters of splenic germinal center (GC) B cells, identified as B220+GL7+CD95+ cells, were assessed by flow cytometry using MitoTracker Red and MitoTracker Green. The ratio of MitoTracker Red to MitoTracker Green fluorescence is shown (n = 7–8 mice per group). **(C)** Unsupervised hierarchical clustering and heatmap of RNA-seq gene-expression profiles from splenic GC B cells isolated at day 8 post-sRBC immunization. Columns represent individual biological replicates and rows represent detected transcripts, with transcript biotypes indicated. Expression values are shown as row-scaled normalized expression. For RNA-seq, n = 3 biological replicates per dietary group, with GC B cells from two spleens pooled to generate each biological replicate. Data in **(B)** are presented as mean ± SEM. Statistical significance was determined by one-way ANOVA followed by Tukey's multiple-comparisons test; exact *P* values are indicated in the figure.

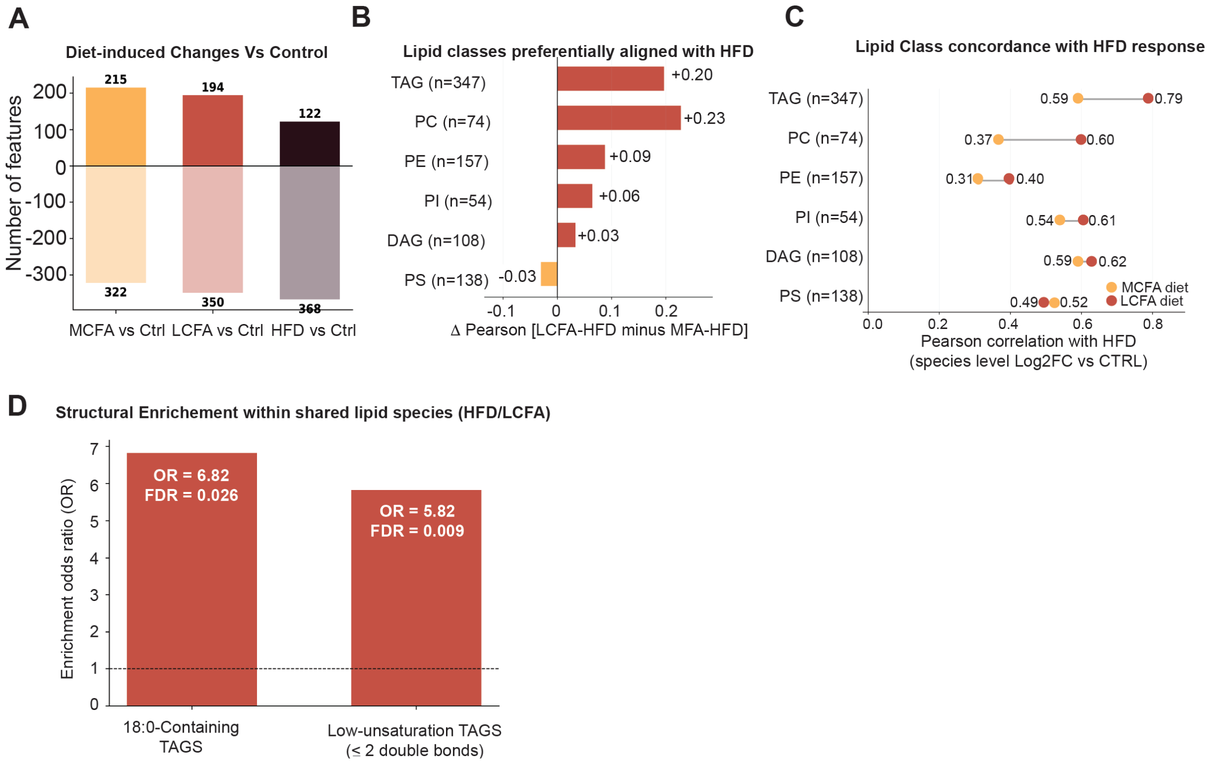


**Figure S4: Circulating lipid remodelling in LCFA-diet and high-fat diet overlap compared to diet rich in medium-chain fatty acids.**

Circulating lipid profiles were analyzed at day 8 following sRBC immunization in mice maintained on control, medium-chain fatty acid (MCFA), long-chain fatty acid (LCFA), or high-fat (HFD) diets. **(A)** Number of circulating lipid features increased (positive values) or decreased (negative values) in each dietary group relative to the control diet. Numbers indicate the total features exhibiting increases or decreases for each comparison. **(B)** Lipid classes preferentially exhibiting concordant responses between the LCFA and HFD diets. For each lipid class, the difference in Pearson correlation with the HFD response between LCFA and MCFA diets (Δ Pearson [LCFA–HFD minus MCFA–HFD]) is shown. Positive values indicate greater concordance of the LCFA response with HFD compared with MCFA. The number of lipid species within each class is indicated. **(C)** Pearson correlations between the HFD-induced lipid response and the corresponding responses to the MCFA or LCFA diets, calculated separately for triacylglycerol (TAG), phosphatidylcholine (PC), phosphatidylethanolamine (PE), phosphatidylinositol (PI), diacylglycerol (DAG), and phosphatidylserine (PS) species. Values represent Pearson correlation coefficients based on species-level log2 fold changes relative to the control diet. **(D)** Structural enrichment analysis among lipid species exhibiting shared responses to the LCFA and HFD diets. Enrichment of TAG species containing stearate

(18:0) and TAG species with low unsaturation (≤2 double bonds) is shown as odds ratios (OR). Statistical significance was assessed by enrichment analysis with correction for multiple testing; false discovery rate (FDR)-adjusted *P* values are indicated. The dashed line denotes an odds ratio of 1 (no enrichment).
